# mosGraphGen: a novel tool to generate multi-omics signaling graphs to facilitate integrative and interpretable graph AI model development

**DOI:** 10.1101/2024.05.15.594360

**Authors:** Heming Zhang, Dekang Cao, Zirui Chen, Xiuyuan Zhang, Yixin Chen, Cole Sessions, Carlos Cruchaga, Philip Payne, Guangfu Li, Michael Province, Fuhai Li

## Abstract

Multi-omics data, i.e., genomics, epigenomics, transcriptomics, proteomics, characterize cellular complex signaling systems from multi-level and multi-view and provide a holistic view of complex cellular signaling pathways. However, it remains challenging to integrate and interpret multi-omics data for mining key disease targets and signaling pathways. Graph AI models have been widely used to analyze graph-structure datasets, and are ideal for integrative multi-omics data analysis because they can naturally integrate and represent multi-omics data as a biologically meaningful multi-level signaling graph and interpret multi-omics data via graph node and edge ranking analysis. However, it is non-trivial for graph-AI model developers to pre-analyze multi-omics data and convert the data into biologically meaningful graphs, which can be directly fed into graph-AI models. To resolve this challenge, we developed **mosGraphGen** (multi-omics signaling graph generator), generating Multi-omics Signaling graphs (mos-graph) of individual samples by mapping multi-omics data onto a biologically meaningful multi-level background signaling network with data normalization by aggregating measurements and aligning to the reference genome. With mosGraphGen, AI model developers can directly apply and evaluate their models using these mos-graphs. In the results, mosGraphGen was used and illustrated using two widely used multi-omics datasets of TCGA and Alzheimer’s disease (AD) samples. The code of mosGraphGen is open-source and publicly available via GitHub: https://github.com/FuhaiLiAiLab/mosGraphGen

## Introduction

Along with the advancements of next-generation sequencing (NGS) and high-throughput technologies, multi-omics datasets including genomics, epigenomics, transcriptomics, proteomics, and metabolomics, have been abundantly generated. The multi-omics datasets characterize the dysfunctional molecular mechanisms of complex diseases from different aspects. The integrative analysis of multi-omics data can provide a holistic view of the mysterious and complex signaling systems of cells from complex diseases. Multi-omics data-driven studies are at the forefront of precision medicine and healthcare, which are critical for uncovering novel therapeutic targets and discovering novel and effective drugs and cocktail treatments. Recently, a new Multi-omics for Health and Disease Consortium was established by the National Institutes of Health (NIH), aiming to advance multi-omics data generation and integrative analysis for human disease and health research.

### Large-scale multi-omics datasets of different diseases have been generated and publicly accessible

For example, multi-omics data of >40,000 cancer samples from ∼69 primary sites were generated in The Cancer Genome Atlas (TCGA)^1,2^ project, which is a comprehensive initiative launched by the National Cancer Institute (NCI) and the National Human Genome Research Institute (NHGRI). The datasets were generated and publicly accessible^1,3^ to characterize and understand the genomic alterations and molecular profiles of >30 cancer types/subtypes. In addition, to increase the range of characterizing the cancer cell lines, the Cancer Cell Line Encyclopedia (CCLE)^4^ study provides multi-omics datasets of ∼1000 cell lines for 36 tumor types. The datasets have been widely used to study genetic biomarkers and associations with drug and cocktail responses. Also, the multi-omics datasets of Alzheimer’s disease (AD) multi-center cohorts e.g., Mayo Clinic^5,6^, Mount Sinai/JJ Peters VA Medical Center Brain Bank (MSBB-AD)^7^, and ROSMAP^8^, have been generated and publicly accessible via the synapse website of AD-AMP^9,10^. Moreover, large-scale multi-omics data, like genetics, epigenetics, transcriptomics, proteomics, and metabolomics^11–13^ are being generated by ongoing NIA-supported exceptional longevity (EL) studies, including the Long-Life Family Study (LLFS), the Longevity Consortium (LC), the longevity genomics (LG), and the Integrative Longevity Omics (ILO), to understand and identify protective factors/targets, biological processes, and signaling pathways that promote health and life span in exceptionally long-lived individuals. These datasets are valuable for studying the complex molecular mechanisms of complex diseases. Moreover, single-level multi-omics datasets are generated. The advance in single cell (sc) omics has made it a powerful technology to investigate the genetic and functional heterogeneity of diverse cell types within disease microenvironment/niche^14,15^. Compared with the tissue-level, single cell (sc) omics datasets provide a finer view of the complex signaling system within diverse cell types such as disease and immune cell types, subtypes, and different cell states^14,15^.

### Multi-omics data integration and interpretation remain an open problem, and network-based models are ideal for multi-omics data analysis

Analyzing and interpreting multi-omics data is complex and computationally challenging. It requires novel and sophisticated bioinformatics and data integration techniques to uncover meaningful insights. The following section summarizes related works of multi-omics data analysis and provides a comprehensive review of existing multi-omics data integration analysis models^16^. Specifically, these models were divided into a few categories: similarity, correlation, Bayesian, multivariate, fusion, and network-based models^16^. Network-based models, such as PARADIGM^17^ (PAthway Representation and Analysis by Direct Inference on Graphical Models), are some of the most widely used methods; Visible Neural Network (VNN^18^) models (e.g., DCell^19^ and DrugCell^20^) were proposed using large hierarchical deep learning architecture to model the hierarchical organization of biological processes and to predict drug response with important biomarkers. However, the signaling cascade level activity has not been specifically investigated in DCell^19^ and DrugCell^20^ models. The recently developed graph neural network (GNN)^21^ model is ideal for graph-structure data analysis tasks and multi-omics data analysis with its ability to 1) integrate multi-omics data as features of nodes and 2) model the signaling interactions or signaling cascades among the proteins using protein-protein interactions^22,23^ or signaling pathways^24,25^; 3) the key signaling target identification and pathway analysis can be achieved via node and edge ranking in GNN^26–30^. Here are a few recent studies that made use of the network-/graph-based analysis for multi-omics data analysis. MOGONET^31^ used the omics-specific similarity graphs among samples and used the GCN^14^ to learn patients’ labels from each omics data independently. The MoGCN^32^ employed a similar design using the patient similarity. GCN-SC^33^ employed GCN to combine single-cell multi-omics data. MOGCL^34^ explored the graph contrastive learning to pretrain the GCN on the multi-omics dataset. However, these models have yet to incorporate signaling pathways.

Computational tools that can generate graph-structure data for individual samples, i.e., mapping multi-omics data onto a biologically meaningful background signaling graph, are urgently needed and critical for developing graph-AI models. As introduced above, we found the top two challenges in graph-AI for multi-omics data analysis are: building a large-scale biologically meaningful background signaling graph to integrate multi-omics data and developing effective approaches to rank key signaling targets and pathways from the large-scale background signaling graph. For many graph-AI developers, particularly those without training in multi-omics data pre-analysis and integrative analysis, it is a challenging task to map the multi-omics data of individual samples onto a meaningful network and generate graphs as inputs ready for graph-AI models. This is because omics-specific pre-analyses are needed to convert the raw data into the standard annotation, like multi-level gene-protein-promoter-enhancer-associations, for multi-omics data integration; and it is essential to link these annotations via biologically meaningful interactomes, including protein-protein interactions, signaling pathways, and transcription factor (TF)-target interactions. To address these challenges, we developed mosGraphGen, a multi-omics signaling graph generator that converts multi-omics datasets into graph-structured data, which can then be used as inputs for graph-AI models. This facilitates integrative and interpretable multi-omics data analysis, such as target and edge ranking analysis within graph-AI models.

#### Open-source

The code of mosGraphGen is open-source and publicly available via GitHub: https://github.com/FuhaiLiAiLab/mosGraphGen. The following sections provide the details of the methodology and results.

## Methodology and Materials

### Multi-omics data pre-processing

**Figure 1** presents a detailed schematic of the pipeline designed to transform extensive multi-omics datasets into graph-based models. This pipeline encapsulates the sequential and methodical approach employed to preprocess, integrate, and ultimately represent data from multi-omics datasets, clinical dataset, reference genome dataset and regulatory network dataset as coherent networks. This pipeline ensures the systematic standardization with aggregating and resolving duplicates across multi-level and integration necessary for generating multi-omics datasets that are critical for advancing our understanding of complex biological systems by referring to the genome dataset. The specific procedures within the pipeline are outlined in the following description.

**Figure 1.**
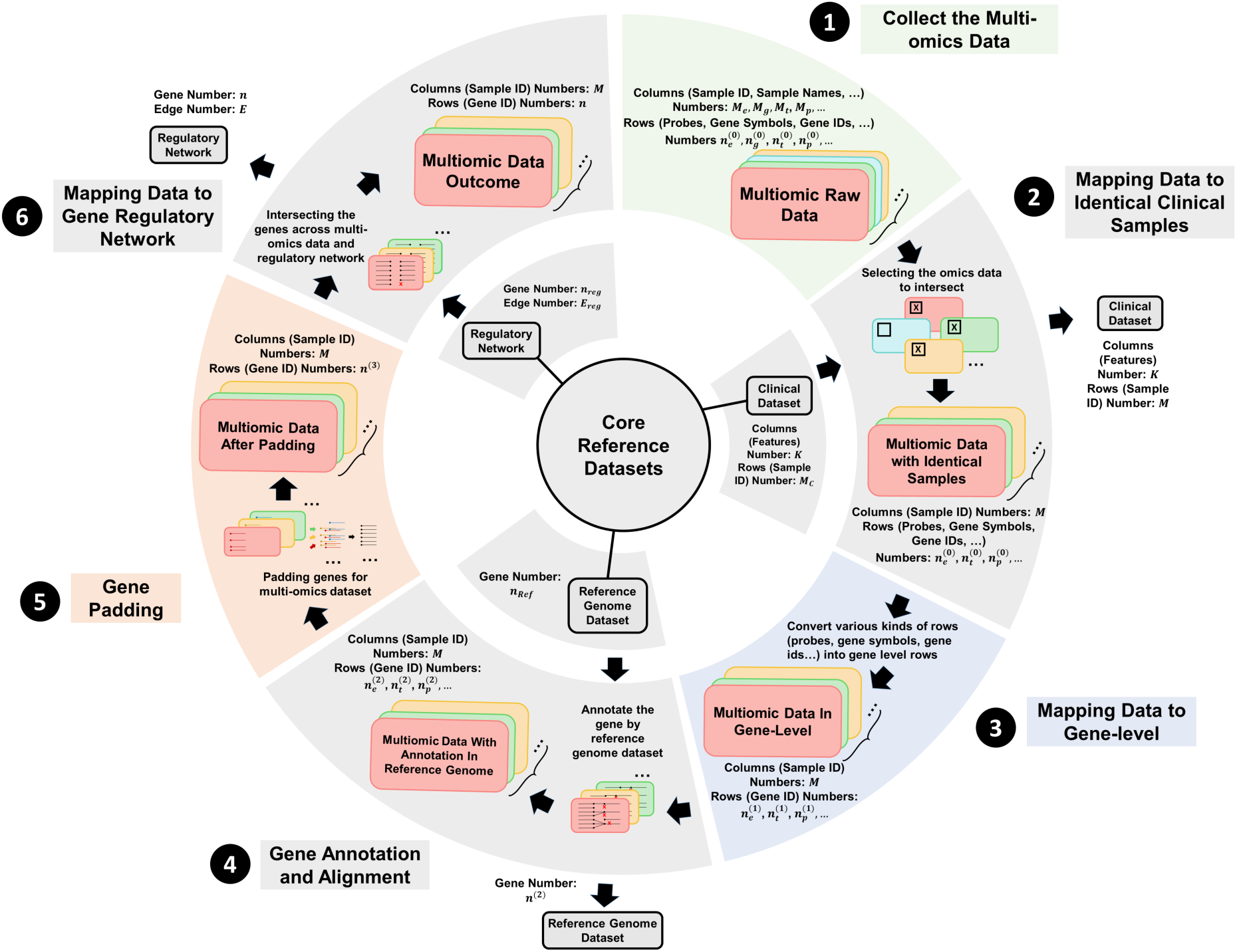
Flowchart multi-omics data processing. ❶Collect various kind of multi-omics data like (Epigenomics, Genomics, Transcriptomics, Proteomics, etc.). ❷Integrating the multi-omics data with clinical data and getting the identical samples across the datasets. ❸Converting the rows (probes, gene symbols, gene ids, etc.) into gene-level by aggregating the same measurements for one gene or by dropping the duplicates for gene synonym. ❹Aligning genes by reference genome so that the final annotation for each gene in multi-omics data will be generated. ❺Unifying the number of genes across multi-omics datasets and make the data imputation by filling zero values in empty spaces. ❻Integrating the gene regulatory network with multi-omics datasets and generating the final multi-omics data with identical number of samples and genes

#### Multi-omics datasets

The multi-omics data can be downloaded from multiple public available datasets. For example, multi-omics data and their related datasets for TCGA dataset and ROSMAP dataset can be downloaded from public available dataset. After downloading the multi-omics (epigenomics, genomics, transcriptomics, proteomics, etc.) datasets from resources, the multi-omics datasets will be converted into 2-dimensional spaces data frames with columns (sample IDs, sample names, etc.) of *M_e_*, *M_g_*, *M_t_*, *M_p_*, … and rows (probes, gene symbols, gene IDs, etc.) of 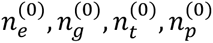, … For example, the methylation data in UCSC dataset have row type of CpG probes target IDs and 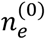 is the number of rows in this methylation dataset. And column type will be sample IDs and *M_e_* is the number of columns in UCSC methylation dataset.

#### Align multi-omics data with clinical information

By selecting the multi-omics datasets from the data downloaded, the clinical dataset with *M_c_* samples and *K* features will be integrated into the multi-omics data. In these clinical datasets, information of each sample (like OS (overall survival), PFI (progression-free interval), DSS (disease-specific survival), DFI (disease-free interval), vital status in UCSC dataset or sex, age_at_visit, ceradsc, cogdx in ROSMAP), will be used as the labels of each sample for different tasks. Since different omics data might use different sample information (for example, OS binary classification value will be used as the label to indicate the survival status for samples in UCSC dataset and ceradsc types will be used as the label to indicate the AD level for each patient in ROSMAP study), the step will associate each omics data to the unique sample ID, which associate with clinical information. Nevertheless, the multi-omics datasets in ROSMAP have the sample ID recognition problem. Hence, the sample ID mapping (e.g. PT-M5AF to R1822146 in ROSMAP dataset) system across multi-omics datasets should be constructed to align identical samples across datasets (check **Table 1** for details).

**Table 1.** Mapping relations example for samples in ROSMAP.

| individualID | mirna_id | mwas_id | mrna_id | cnvdata_id | projid | Study |
| --- | --- | --- | --- | --- | --- | --- |
| R1822146 | b01.127N | PT-M5AF | 575_120521_2 | SM-CTDQQ | 20271359 | ROS |
| R3143439 | b01.128N | PT-BZBT | 502_120515_1 | SM-CTEMJ | 38967303 | MAP |
| R6879714 | b01.130C | PT-3PTN | 660_120530_1 | SM-CTEEU | 65736039 | MAP |
| R8963331 | b01.130N | PT-M5GT | 607_120523_2 | SM-CJEKL | 20197364 | ROS |
| ... | ... | .. | ... | ... | ... | ... |

However, only a part of sample IDs in different omics data are overlapped, which will reduce the *M_c_* samples to identical number of samples with *M* in the selected datasets (e.g., epigenomics, transcriptomics proteomics, etc.), depending on the omics data selection strategies. Also, after the intersection with the clinical datasets, the number of rows in selected datasets (e.g., epigenomics, transcriptomics proteomics, etc.) will be 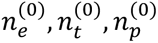,etc.

#### Mapping data to gene-level

Given that the rows of each multi-omics dataset vary, the methods used for mapping the rows to gene level are unique for each dataset. In details, each omics data have their special mapping steps described as follows:

##### (1) Epigenomics/methylation data pre-analysis

The methylation data will be retrieved from the UCSC XENA DNA methylation (450k) for UCSC dataset and ROSMAP_arrayMethylation_imputed for ROSMAP dataset. However, the rows in above methylation datasets are probes to detect the methylation condition in each individual CpG site. To convert the CpG site level methylation to TSS level, the GEO GPL16304 Platform will be considered for both datasets, which provides annotations, such as Distance_closest_TSS and Closest_TSS_gene_name, for each probe in the methylation (450k) dataset, mapping probe IDs to their corresponding gene names (e.g., cg00001583 to NR5A2). Furthermore, the gene name from the GPL16304 file was replaced with the probe ID in the methylation data by merging with the probe ID file. Hence, the nearest TSS gene name was incorporated into the new data frame.

In the next step, a preliminary analysis is essential to convert the TSS level methylation to gene level, focusing specifically on the −6 kb to +3 kb range around transcription start sites (TSS). This range will be subdivided into five distinct regions and probes located outside this specified range will be excluded from the analysis, which will convert rows from the CpG site level to gene-level^35–39^. In detail, promoter regions were delineated into three categories: the core promoter region (0-50bp upstream of TSS), the proximal promoter region (50-250bp upstream of TSS), and the distal promoter region (250-3000bp upstream of TSS). Additionally, the area 3000 to 6000 bp upstream of the distal promoter region was defined as the upstream region, while the downstream region encompassed the area from the TSS extending to 3000 bp downstream (check **Figure 2** for division of promotor and whole methylation data regions).

**Figure 2.**
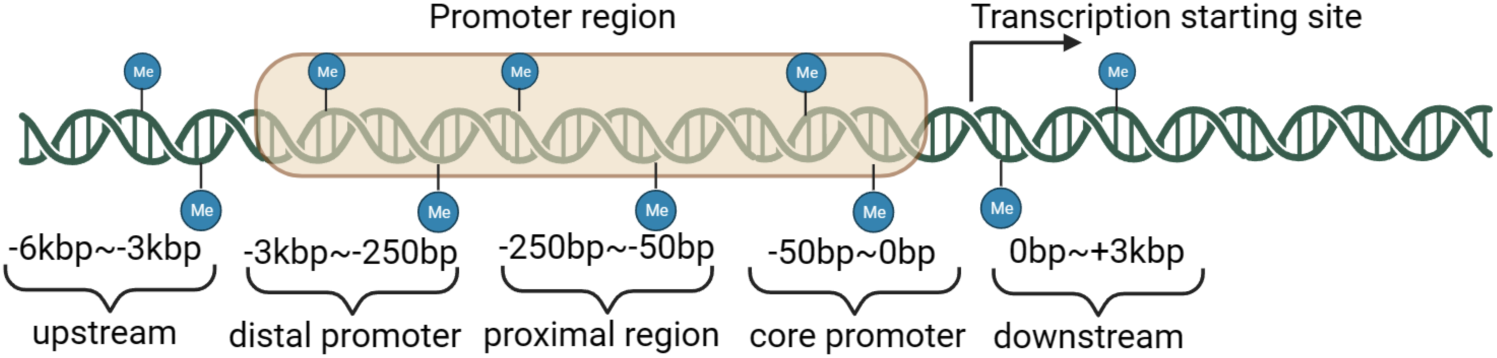
Dividing methylation regions into 5 regions

According to the defined regions of interest for methylation values, the TSS level methylation will be transformed into the gene level by using mean aggregation and rows with TSS closest distances outside above regions will be omitted. This resulted in five CSV files, each containing the average methylation levels for genes in distinct 5 regions as mentioned before. To keep the dimensions of methylation data in each regions in same format, the unique genes in all the regions should be integrated and padding the empty value with zero and the number of rows in methylation data will be 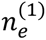 for UCSC or ROSMAP dataset.

##### (2) Genetic variations pre-analysis

For UCSC genome variant datasets, the preprocessed genomic dataset is provided in gene-level mutation with binary value (0 wild-type for and 1 for non-silent mutation). To remove the duplicated rows named with same gene, mean values was applied to aggregate the mutations. With respect to Copy Number Variations data in ROSMAP dataset, we employed the pyensembl package to assign gene names based on the chromosomal start and end positions (e.g., Start-End 830676-834492 to gene LINC01128). Given multiple types of variation are defined as deletions (DEL), duplications (DUP), and multiple copy number variations (mCNV), genome variants data were summed based on multiple genetic variation by variation types to capture the cumulative effect of variations of the same gene. In contrast to other multi-omics data, to keep the dimensions of genetic variation data in each type of variation in same format, the unique genes in all the variation types should be integrated and padding the empty value with −1 to maintain the integrity of the variation dataset. In the end, the output dataset after this processing steps will have the number of rows as 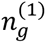 for UCSC or ROSMAP dataset.

##### (3) Transcriptomics data pre-analysis

For UCSC RNA sequencing datasets, gene names were directly converted using ID mapping table associated with the gene expression dataset from UCSC website. With regard to ROSMAP RNA sequencing data, we removed the version suffix from gene IDs in the RNA sequencing data (e.g., transforming ENSG00000167578.11 to ENSG00000167578) to adapt version updates in annotations. Using the official Ensembl dataset as the reference, our sequencing data was refined by replacing gene IDs with their respective gene names based on the established correspondence between gene IDs and names within the Ensembl resource. Mean values were calculated for duplicate gene symbols to ensure a single, representative value for each gene for both UCSC and ROSMAP transcriptomics dataset, resulting the number of rows as 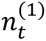.

##### (4) Proteomic data pre-analysis

Regarding proteomics data in ROSMAP dataset, we first discarded protein accession numbers, focusing solely on gene/protein names (e.g., VAMP1|P23763 to VAMP1) to ensure consistency with the gene-centric analysis framework of mosGraphGen. In contrast to other omics datasets, proteomics data in both UCSC and ROSMAP datasets did not require a specific pre-analysis and mapping phase due to its direct association with gene/protein names. Mean values were calculated for duplicated genes to ensure a single, representative value for each gene, ending with the number of rows as 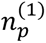.

#### Gene annotations, alignment and padding

After getting the gene-level data for selected multi-omics datasets, the reference genome dataset (e.g. Ensembl dataset, containing *n_ref_* genes) will be chosen as the standard for annotating the genes across multi-omics datasets. Therefore, a comprehensive comparison was conducted with the Ensembl dataset to ensure the coherence and accuracy of our genome variant dataset. Genes identified in pre-processed multi-omics datasets that were not present in the Ensembl dataset were excluded from further analysis to ensure that only genes with recognized genomic annotations were included. Therefore, the selected multi-omics datasets will contain *M* identical samples and rows with 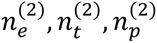, … To unify the number of genes across multi-omics datasets, padding methods will be applied here with integrating unique genes in all datasets. Hence, zero values will be filled in the empty spaces to complete this data imputation method. Specifically, genomics data will be filled with −1 for empty spaces to distinguish the empty values with the non-mutation conditions. After this padding step, the number of genes will be identical across all multi-omics datasets with *n*^(3)^ rows.

#### Mapping data to gene regulatory network

To integrate the gene regulatory network for multi-omics data, the selected gene regulatory network (e.g. KEGG^24,40^, BioGRID^23,41^ or STRING^42^) with *n_reg_* genes and *E_reg_* protein-protein interactions will be intersected with the multi-omics datasets. Filtering out the genes not overlapped with the gene regulatory network, the multi-omics datasets will contain the identical number of samples *M* and genes *n*.

## Results

### Overview of mosGraphGen

From a computational perspective, integrating multi-omics data into the graph model involved several steps. First, multi-omics data from each patient were collected. In the meanwhile, the related datasets such as clinical datasets as sample labels, reference genome dataset as gene annotation and gene regulatory network as organizing the gene knowledge graph will also downloaded and integrated into multi-omics dataset as mentioned in the method part of multi-omics data pre-processing. However, most of the networks ignored the details of biological processes such as methylation and transcription. Hence, the detailed version was depicted in **Figure 3** with organizing the promoter regions and transcription activity. Therefore, the re-organized network could be leveraged as the universal knowledge graph for each patient as the graph model. The input features for each individual patient were generated by integrating the matrices from each omics feature. Utilizing the network and feature inputs, graph neural network models were trained to predict patient outcomes and identify patient-specific critical biomarkers. This was achieved by extracting attention scores or edge weights from the trained models. The complete processes, including data loading, model training, and downstream analysis, is publicly accessible on GitHub documents: https://github.com/FuhaiLiAiLab/mosGraphGen/blob/main/README.md

**Figure 3.**
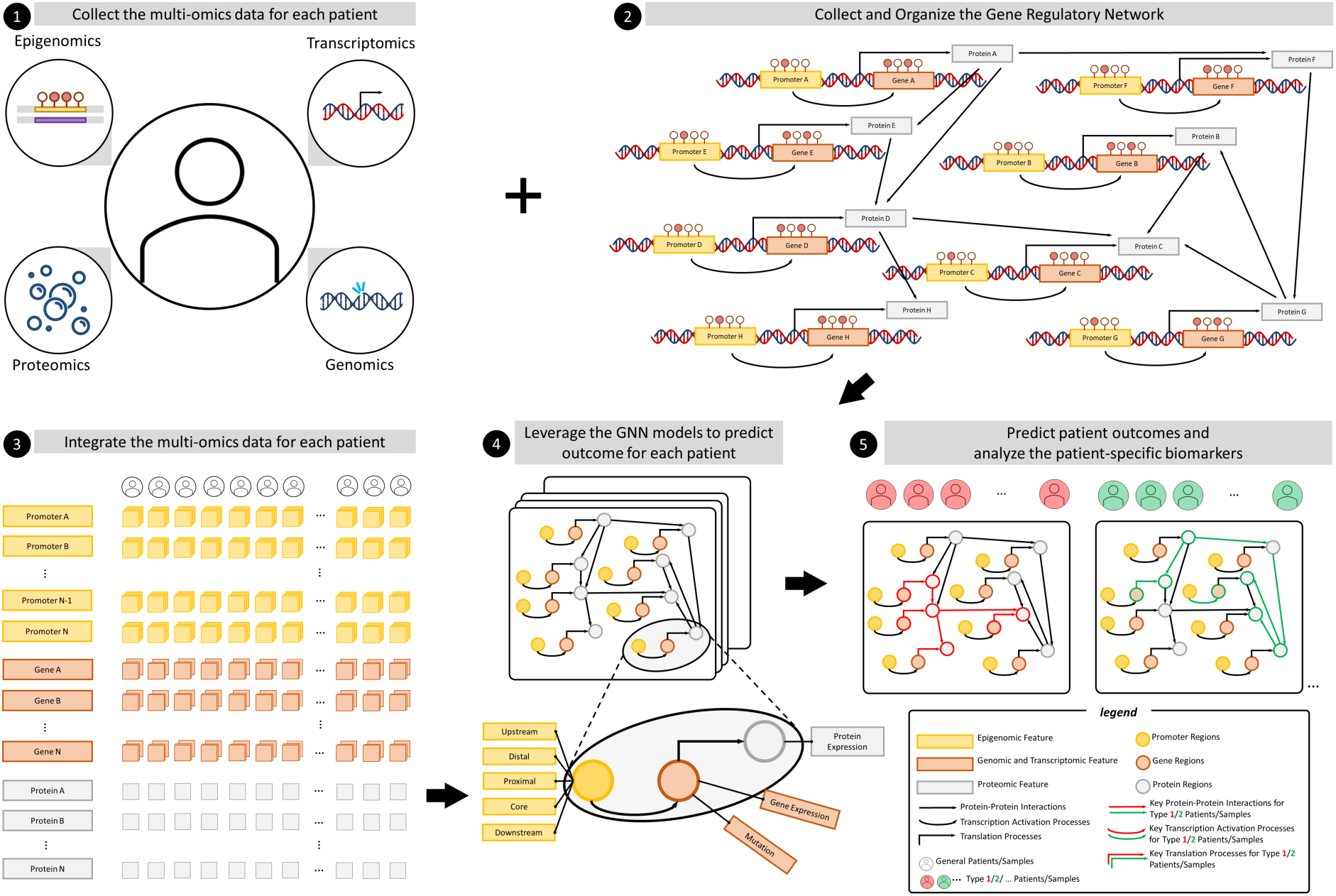
Overview of mosGraphGen. ❶Multi-omics dataset (Epigenomics, Transcriptomics, Genomics and Proteomics) for each individual patient will be collected. ❷The gene regulatory network from KEGG, BioGRID or STRING will be downloaded and integrate the protein-protein interactions with the promoters and transcriptions process. Hence, the re-organized regulatory network will incorporate more details and be formalized as the input matrix for graph model. ❸The collected multi-omics dataset will be processed and integrate into the matrices format for each individual patient. ❹The well-formatted matrices (features and regulatory network) will be used as the input for various graph neural network models to make the prediction for each individual patient. ❺Downstream analysis will be made by extracting the edge weights from the attention trained from the selected graph neural network. Hence, the patient-specific analysis of discovering the critical biomarkers will be generated.

### Graphical overview of mosGraphGen

**Figure 3** shows the architecture of the graph neural network model. The model has input 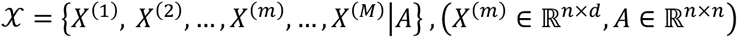, where *M* denotes the number of samples/patients in each input dataset, and *n* denotes the number of nodes/genes used in the graph AI model; *A* is adjacency matrix for number of *n* nodes for the generated multi-omics signaling graph; and *d* is the number of multi-omics features organized in the initial embeddings in each node. To predict the outcome of all samples 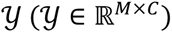, where *C* is the number of classes and 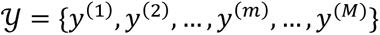, we build a machine learning model *f*(⋅) with 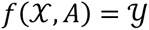, where *X* denotes all of the data points in the dataset.

#### Graph-AI ready data format

For both UCSC and ROSMAP datasets, the processed datasets were saved in the special folder in CSV format firstly. Afterwards, the clinical datasets were considered as the graph prediction labels, i.e., the 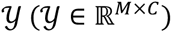 in the methodology part. Another task was to combine the multi-omics features in each individual CSV file by concatenating those features into a 3D NumPy array, i.e., *X* in the math part, which contained information about each multi-omics feature in all genes for every sample/patient. Since the torch-geometric package was used here to enhance the computing speed and save more spaces in GPU, the adjacency matrix was transformed into the edge index format with the dimension of (*E*, 2), where *E* was the number of edges in the proposed graph model or the number of edges/links in the gene regulatory network. In addition, to validate the proposed model, a 5-fold cross-validation was used. In total, there were 3592 model input data points for the UCSC dataset and 138 model input data points for the ROSMAP dataset if all multi-omics datasets are selected. In each fold model, 4 folds of the data points were used as the training dataset, and the rest 1 fold was used as the testing dataset. Finally, the processed data with NumPy format 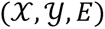 were loaded with DataLoader class in torch-geometric to transform them into the machine readable CUDA tensor (check **Figure 4** for details).

**Figure 4.**
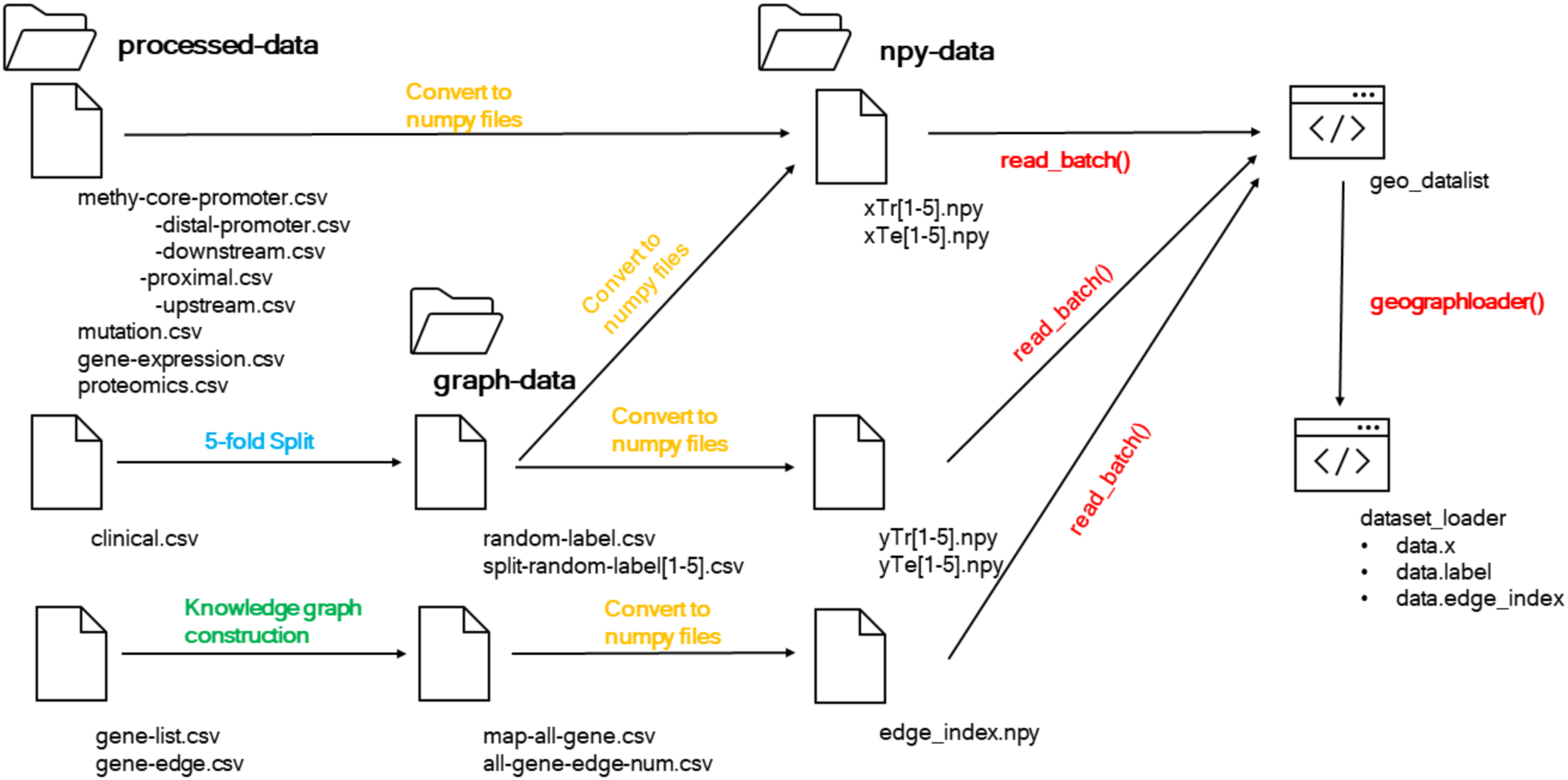
Loading processed data into graph-AI format

### Multi-omics datasets of TCGA cancer samples

**Table 2** presents download links for multi-omics datasets of TCGA cancer samples, encompassing analyses of DNA methylation patterns, somatic mutations—including Single Nucleotide Polymorphisms (SNPs) and Insertions/Deletions (INDELs) — as well as gene expression profiles obtained through RNA sequencing. As to methylation data, the conversion from CpG site-level data to gene-level data will be conducted based on the aforementioned method. The detailed data processing results are described as follows.

**Table 2.**
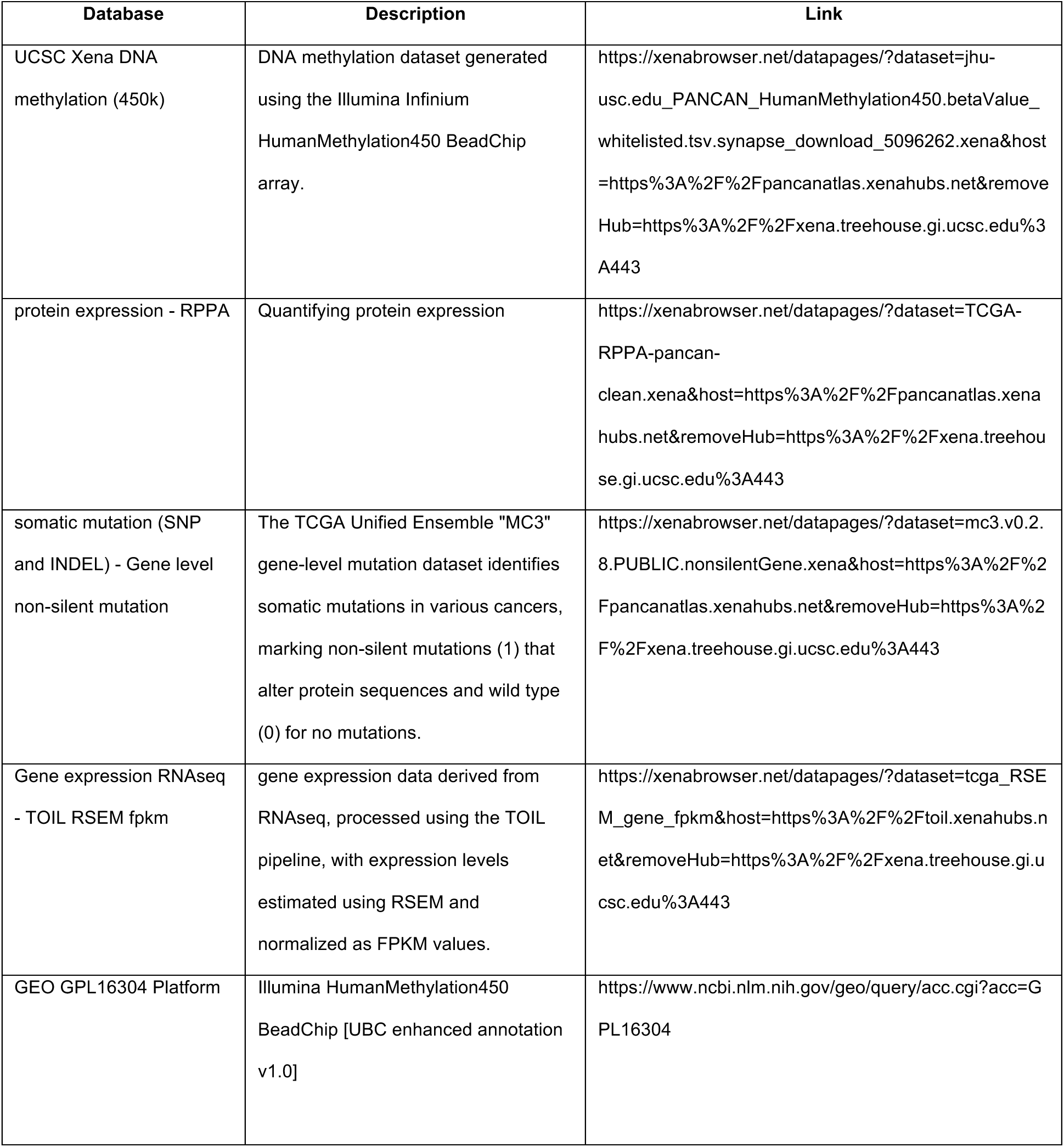
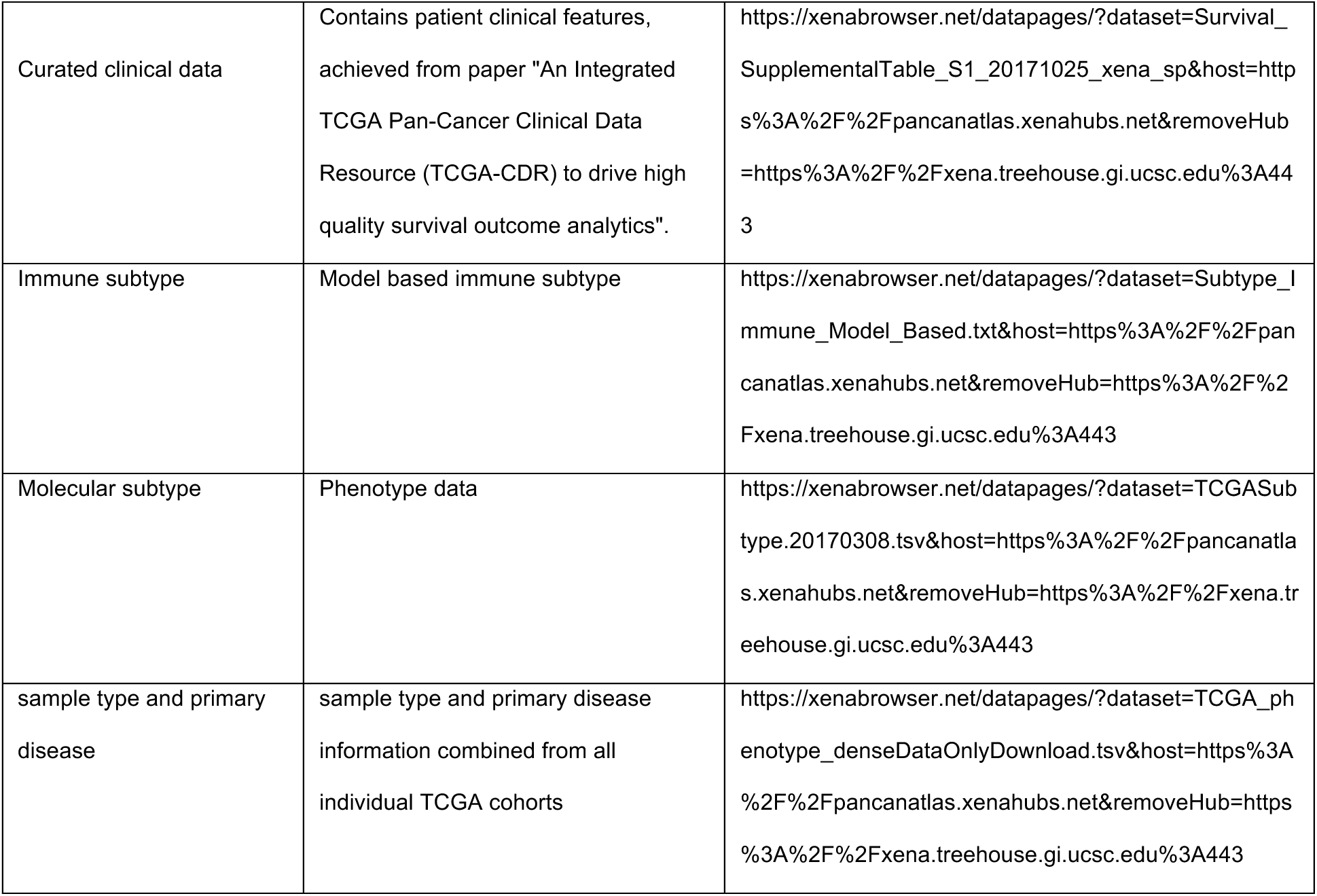
UCSC Database resources.

| Database | Description | Link |
| --- | --- | --- |
| UCSC Xena DNA methylation (450k) | DNA methylation dataset generated using the Illumina Infinium HumanMethylation450 BeadChip array. | https://xenabrowser.net/datapages/?dataset=jhu-usc.edu_PANCAN_HumanMethylation450.betaValue_whitelisted.tsv.synapse_download_5096262.xena&host=https%3A%2F%2Fpancanatlas.xenahubs.net&removeHub=https%3A%2F%2Fxena.treehouse.gi.ucsc.edu%3A443 |
| protein expression - RPPA | Quantifying protein expression | https://xenabrowser.net/datapages/?dataset=TCGA-RPPA-pancan-clean.xena&host=https%3A%2F%2Fpancanatlas.xenahubs.net&removeHub=https%3A%2F%2Fxena.treehouse.gi.ucsc.edu%3A443 |
| somatic mutation (SNP and INDEL) - Gene level non-silent mutation | The TCGA Unified Ensemble "MC3" gene-level mutation dataset identifies somatic mutations in various cancers, marking non-silent mutations (1) that alter protein sequences and wild type (0) for no mutations. | https://xenabrowser.net/datapages/?dataset=mc3.v0.2.8.PUBLIC.nonsilentGene.xena&host=https%3A%2F%2Fpancanatlas.xenahubs.net&removeHub=https%3A%2F%2Fxena.treehouse.gi.ucsc.edu%3A443 |
| Gene expression RNAseq - TOIL RSEM fpkm | gene expression data derived from RNAseq, processed using the TOIL pipeline, with expression levels estimated using RSEM and normalized as FPKM values. | https://xenabrowser.net/datapages/?dataset=tcga_RSEM_gene_fpkm&host=https%3A%2F%2Ftoil.xenahubs.net&removeHub=https%3A%2F%2Fxena.treehouse.gi.ucsc.edu%3A443 |
| GEO GPL16304 Platform | Illumina HumanMethylation450 BeadChip [UBC enhanced annotation v1.0] | https://www.ncbi.nlm.nih.gov/geo/query/acc.cgi?acc=GPL16304 |
| Curated clinical data | Contains patient clinical features, achieved from paper "An Integrated TCGA Pan-Cancer Clinical Data Resource (TCGA-CDR) to drive high quality survival outcome analytics". | https://xenabrowser.net/datapages/?dataset=Survival_SupplementalTable_S1_20171025_xena_sp&host=https%3A%2F%2Fpancanatlas.xenahubs.net&removeHub=https%3A%2F%2Fxena.treehouse.gi.ucsc.edu%3A443 |
| Immune subtype | Model based immune subtype | https://xenabrowser.net/datapages/?dataset=Subtype_Immune_Model_Based.txt&host=https%3A%2F%2Fpancanatlas.xenahubs.net&removeHub=https%3A%2F%2Fxena.treehouse.gi.ucsc.edu%3A443 |
| Molecular subtype | Phenotype data | https://xenabrowser.net/datapages/?dataset=TCGASubtype.20170308.tsv&host=https%3A%2F%2Fpancanatlas.xenahubs.net&removeHub=https%3A%2F%2Fxena.treehouse.gi.ucsc.edu%3A443 |
| sample type and primary disease | sample type and primary disease information combined from all individual TCGA cohorts | https://xenabrowser.net/datapages/?dataset=TCGA_phenotype_denseDataOnlyDownload.tsv&host=https%3A%2F%2Fpancanatlas.xenahubs.net&removeHub=https%3A%2F%2Fxena.treehouse.gi.ucsc.edu%3A443 |

As detailed in method above, raw multi-omics datasets in UCSC are collected and loaded into pandas dataframe (check Table 3 for rows and columns information). The methylation dataset encompasses 396065 probes, targeting the CpG sites of genes across. Since the regions of interest around TSS were predefined as mentioned in **Figure 2**, rows with TSS distances that fell outside these areas were excluded and each CpG probe was assigned with corresponding TSS closest gene name, reducing the number of rows from 396065 to 242313. By splitting the methylation files into 5 files according to predefined promoter regions, the number of rows (genes) will be 5638, 17797, 16169, 10139, and 21757 for the upstream region, distal promoter region, proximal promoter region, core promoter region, and downstream promoter region. After the unique genes in all the regions were integrated and padded, the final number of rows were 24396 for all 5 files.

**Table 3.**
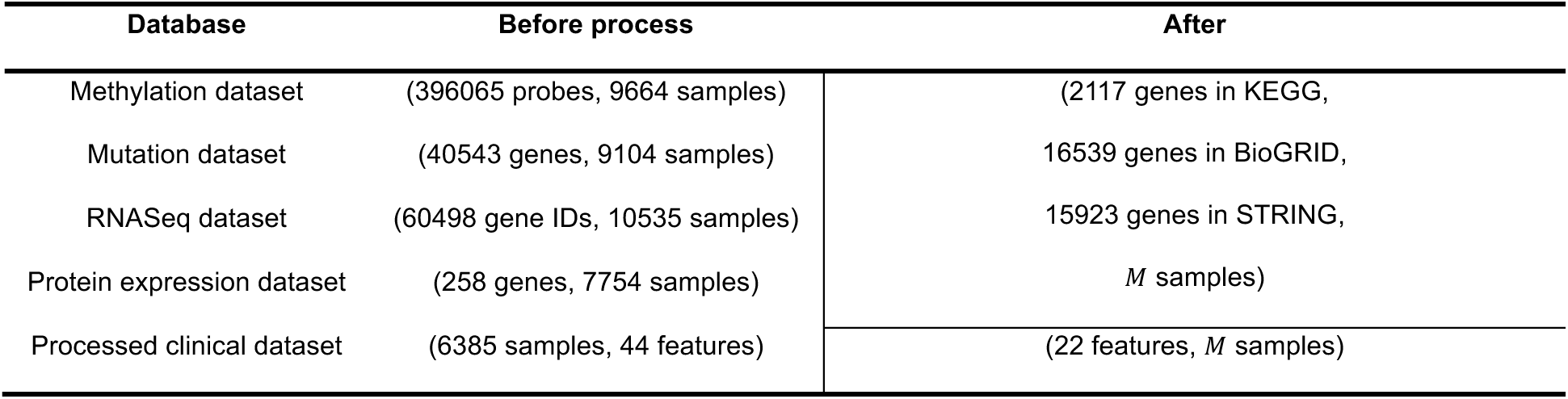
Dimensions of the UCSC multi-omics data before and after the data processing.

For UCSC genome variant datasets, values of duplicated genes were averaged as described in method part to represent the effect of variations of the same gene which reducing the number of rows from 40543 to 40490. With respect to UCSC RNA sequencing datasets, mean values were calculated for duplicate genes to ensure a single, representative value for each gene, resulting number of rows from 60498 to 49432. The UCSC protein expression was also checked with duplicated gene names problem and found no duplication. Therefore, the number of rows (proteins) remains as initial number as 258 (check **Figure 5** for details).

**Figure 5.**
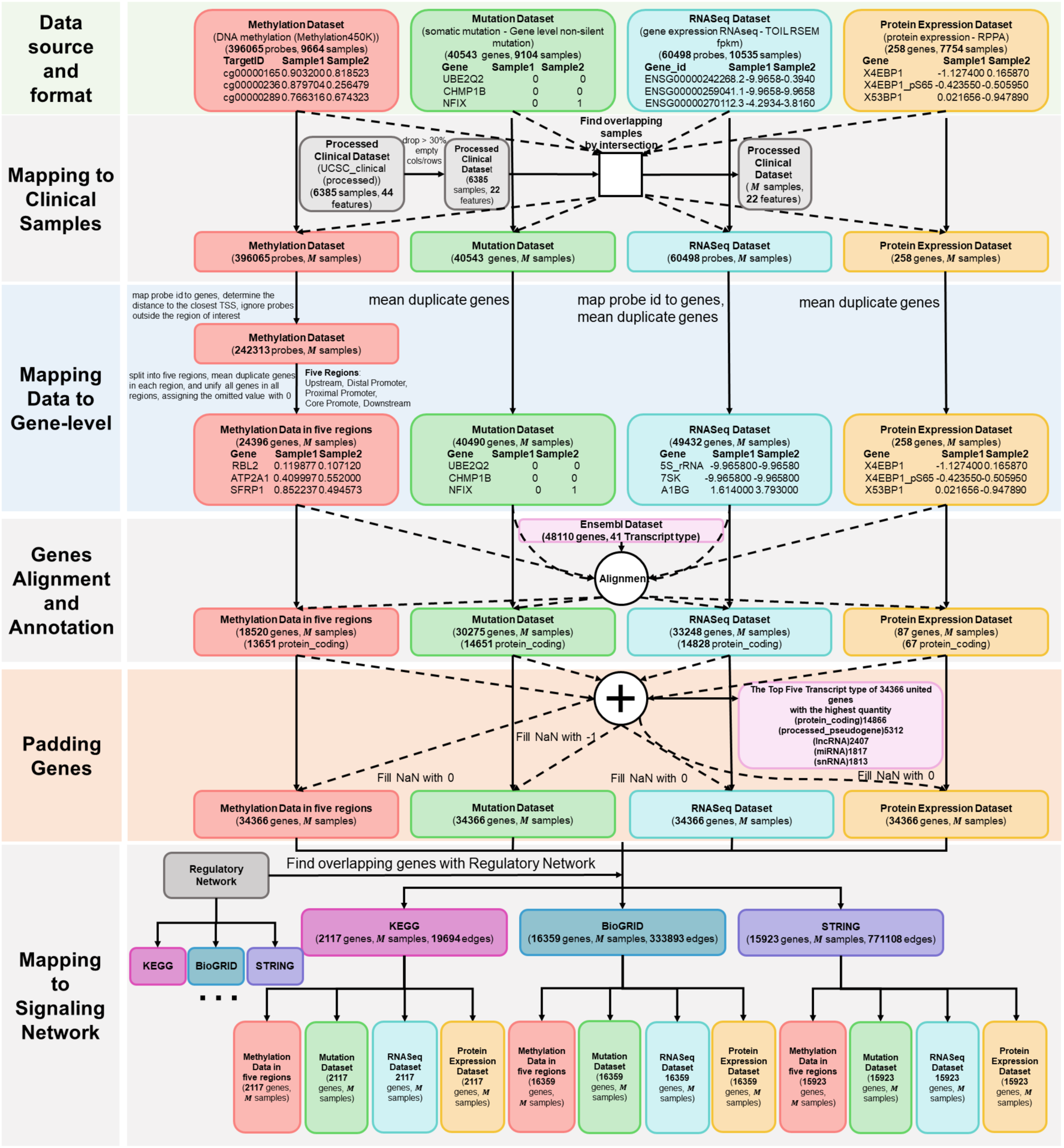
Flowchart of TCGA cancer multi-omics data analysis. (a) The Methylation Dataset undergoes probe-to-gene mapping, with a focus on the −6 kb to +3 kb range around the TSS, followed by division into five genomic regions. (b) The Protein Expression Dataset includes 258 genes. (c) The RNAseq Dataset consists of 60498 probes. (d) The Mutation Dataset is unified by gene identifiers across 40543 genes. These datasets are then integrated to form a unified gene set. (e) Duplicate genes in each dataset are unified by taking mean values (f) All the datasets are filtered using Ensembl Dataset. (g)Missing values are imputed with zeros or negative ones, as appropriate. (h) Integration with the KEGG database refines the data to 2117 common genes. (i) Clinical dataset preprocessing involves dropping or imputing missing values, leading to *M* common samples for multi-omic analysis.

Furthermore, the reference datasets (clinical features, Ensembl dataset, gene regulatory network) were collected and integrated with processed multi-omics dataset for providing labels, gene annotation and knowledge graph construction). Firstly, an examination of four clinical feature datasets sourced from the UCSC databases was conducted. The concatenation of these datasets yielded a composite dataset encompassing 6,385 samples and 22 features, following the exclusion of any features exhibiting greater than 30% missing values. In multi-omics dataset, the number of samples are 9664, 9104, 10535 and 258 in methylation, mutation, RNASeq and protein expression dataset. Afterwards, the common samples/patients across multi-omics datasets and concatenated clinical dataset were intersected, leaving *M* common samples, where *M* = 3592 if all of the multi-omics datasets were utilized (see **Figure 6A** and **Table 3**).

**Figure 6.**
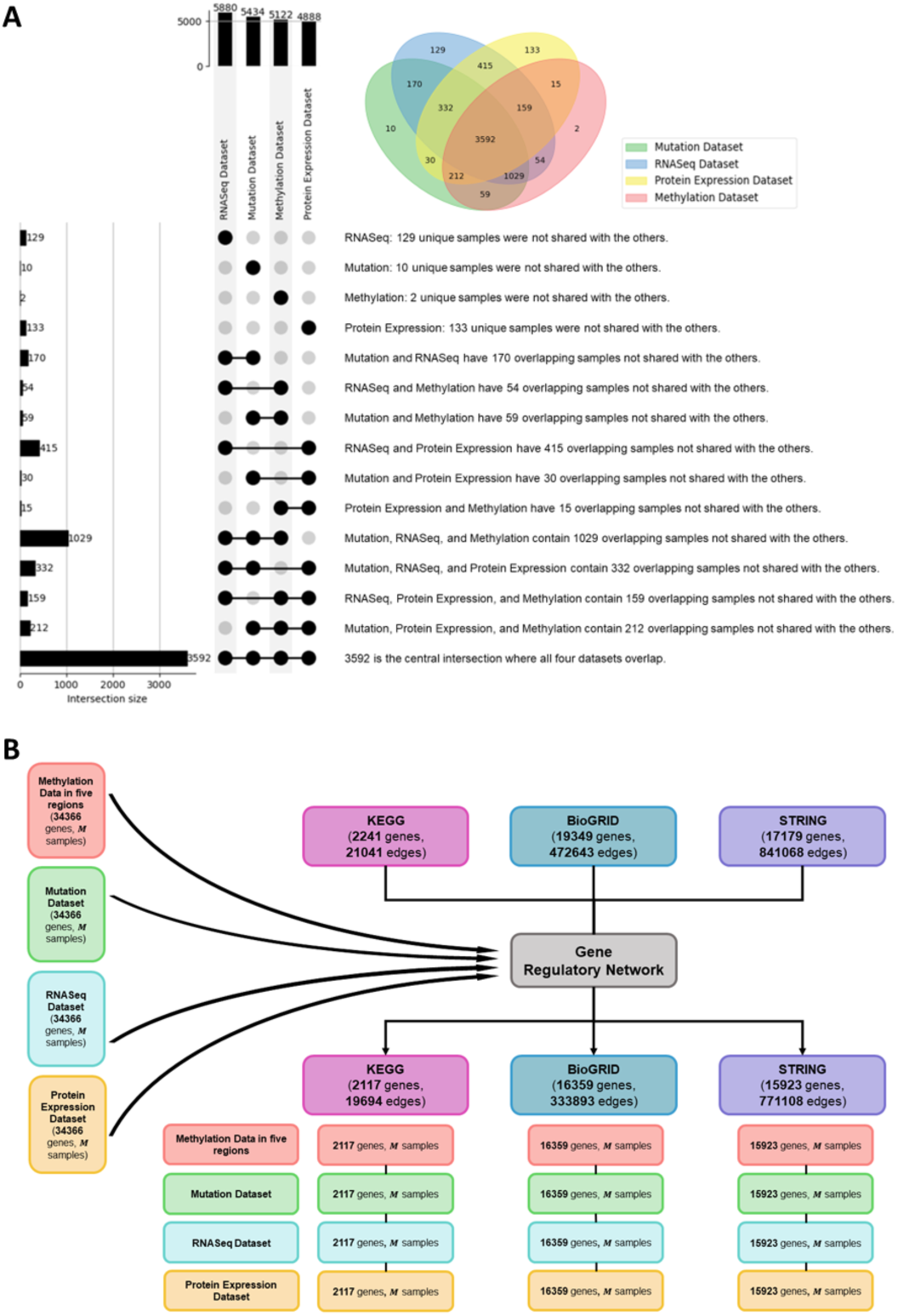
Details for intersection for samples gene regulatory network. (a)The intersection of omics data with clinical samples, finally *M* samples across all omics data. (b) The intersection of the multi-omic data with different options of gene regulatory network

What’s more, Ensembl dataset was leveraged to annotate gene in multi-omics data, resulting the dimensions for these files were methylation dataset (18520 genes, *M* samples), protein expression dataset (87 genes, *M* samples), RNASeq dataset (33248 genes, *M* samples), and mutation dataset (30275 genes, *M* samples). To obtain the union of genes from multi-omics data, all unique genes were collected across the datasets, resulting in a total of 34,366 genes. Then, the genes used to construct the knowledge graph were collected by intersecting genes in multi-omics datasets and gene regulatory network (KEGG^24,40^ (2241 genes, 21041 edges), BioGRID^23,41^ (19349 genes, 472643 edges) and STRING^42^ (17179 genes, 841068 edges)), ending with the number of gene entities as 2117, 16359 and 15923 (see **Figure 6B**).

### Multi-omics datasets of Alzheimer’s Disease (AD)

**Table 3** presents the multi-omics datasets and the corresponding dataset’s download links. The mutation data, derived from “Whole genome sequencing–-based copy number variations reveal novel pathways and targets in Alzheimer’s disease”, were categorized into three distinct mutation types. For the RNA-seq data, null values will be replaced with 0 to ensure the data is meaningful and suitable for modeling purposes. A crucial step in the preprocessing stage was the unification of genes across each dataset. Once all processes were completed, all datasets were collectively processed. The detailed pre-analysis pipeline is described as follows.

Described in method aforementioned, raw multi-omics datasets in ROSMAP are collected and loaded into pandas dataframe (check **Table 4** for rows and columns information). The methylation dataset encompasses 420132 probes, targeting the CpG sites of genes. Since the regions of interest around TSS were predefined as mentioned in **Figure 2**, rows with TSS distances that fell outside these areas were excluded and each CpG probe was assigned with corresponding TSS closest gene name reducing the number of rows from 420132 to 249506. By splitting the methylation files into 5 files according to predefined promoter regions, the number of rows (genes) will be 15812, 20577, 18238, 9871, and 7128 for the upstream region, distal promoter region, proximal promoter region, core promoter region, and downstream promoter region. After the unique genes in all the regions were integrated and padded, the final number of rows were 23280 for all 5 files.

**Table 4.**
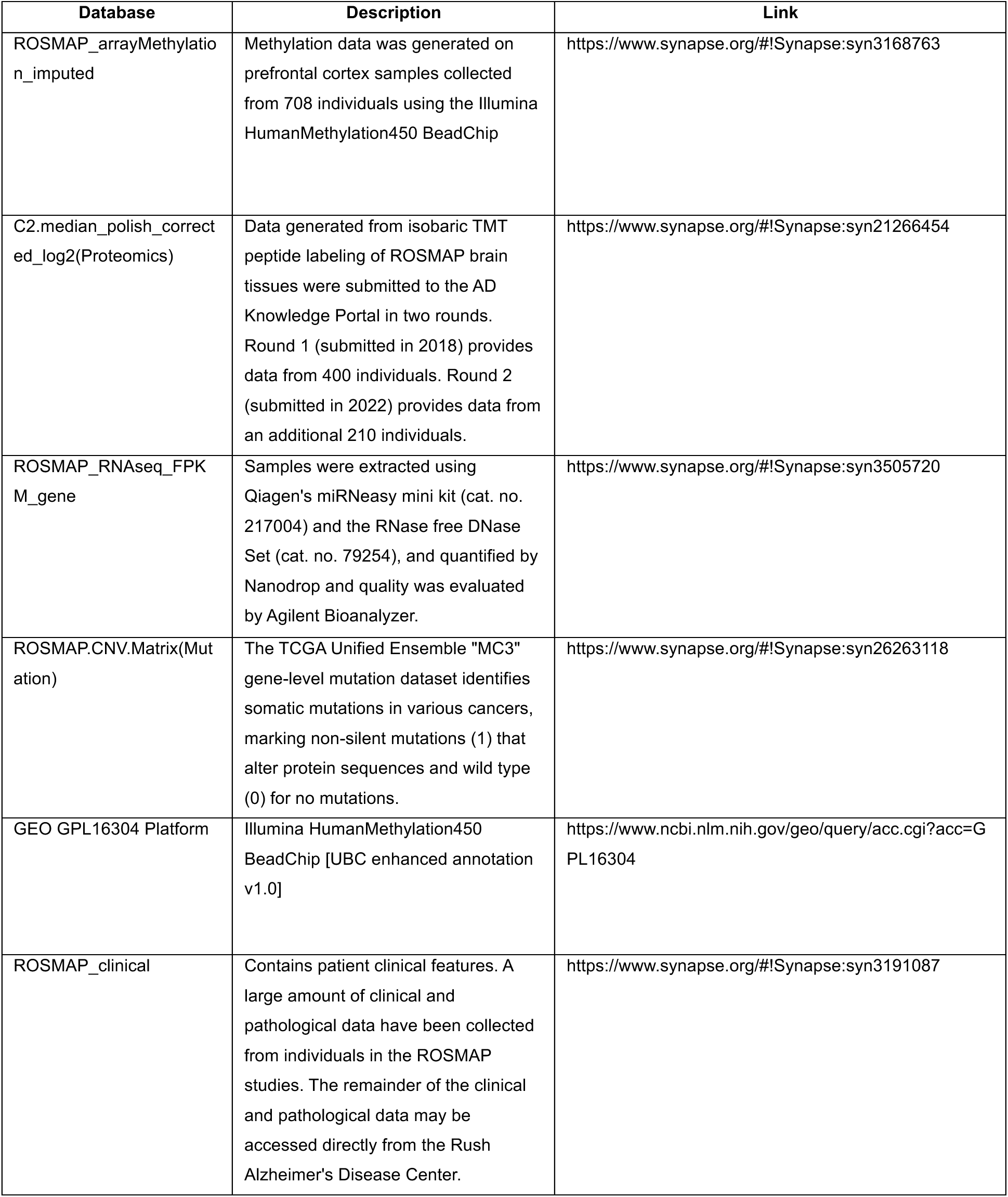
ROSMAP Database resources.

With respect to Copy Number Variations dataset, data were aggregated with summation for duplicate genes to capture the effect of variations across multiple instances of the same gene. To split and unify data, genome variants data were categorized based on the nature of the variation—deletions (DEL), duplications (DUP), and multiple copy number variations (mCNV). And there were three final aggregated files for ‘DEL’, ‘DUP’, and ‘mCNV’ with number of rows with 2483, 1096 and 3726. By identifying and adding all unique gene names to each file, the three files had the same format, generating 6423 unique genes for all 3 types of variation. For any duplicate genes in ROSMAP RNA sequencing data, mean values were calculated to ensure a single, representative value for each gene, resulting number of rows from 55889 to 37898. The proteomics in the ROSMAP dataset was also examined for the problem of duplicated gene names, and gene uniqueness was ensured by merging duplicates, reducing the gene count from 8817 to 8252 (check **Figure 7** for details).

**Figure 7.**
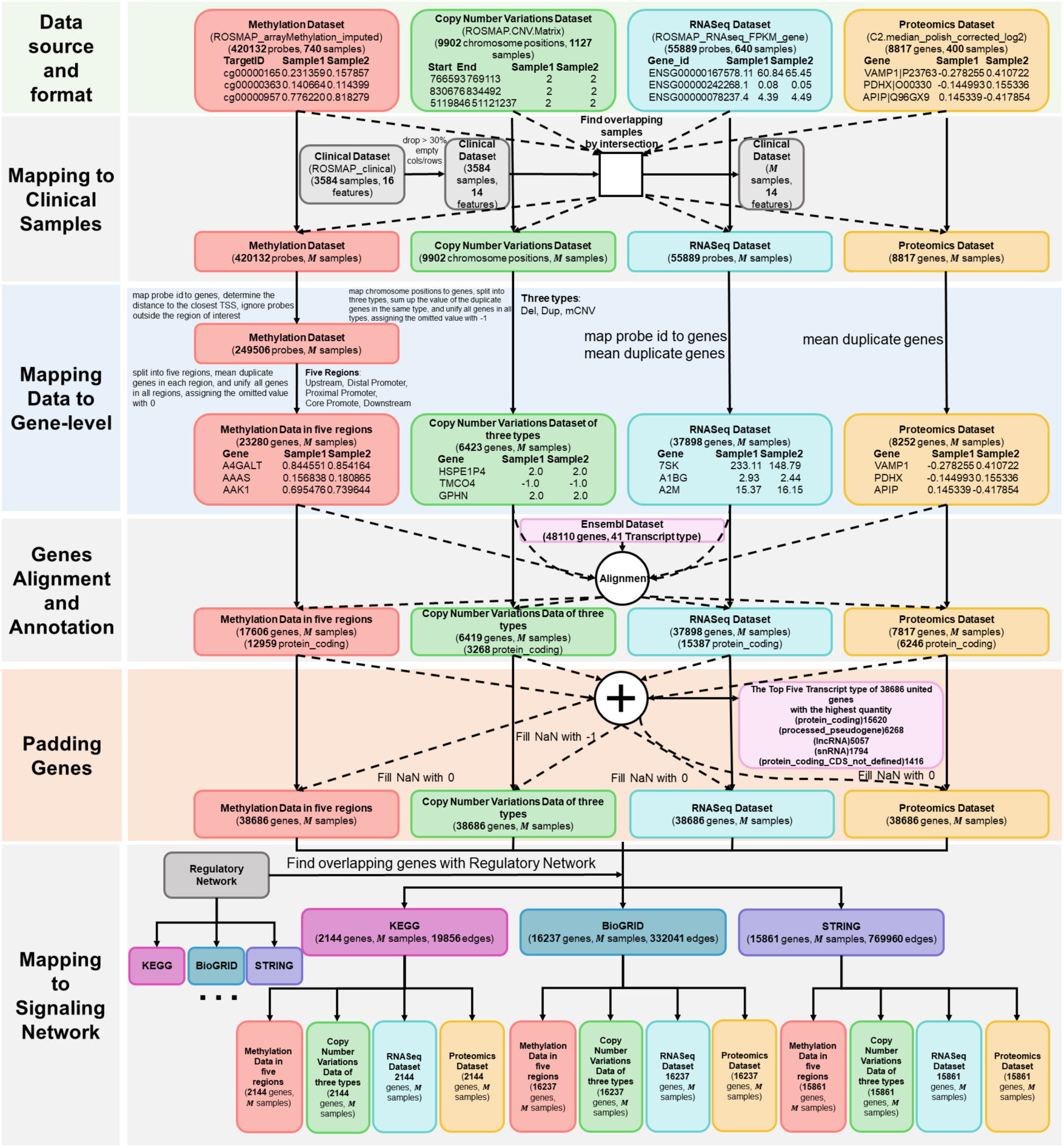
Flowchart of AD multi-omic data processing. (a) The Methylation Dataset undergoes probe-to-gene mapping, with a focus on the −6 kb to +3 kb range around the TSS, followed by division into five genomic regions. (b) The Proteomics Dataset includes 8817 genes. (c) The RNAseq Dataset consists of 55889 probes (d) The Copy Number Variations Dataset undergoes chromosome position-to-gene mapping, followed by division into three types. These datasets are then integrated to form a unified gene set. (e) Duplicate genes in each dataset are unified by taking mean values (f) All the datasets are filtered using Ensembl Dataset. (g) Missing values are imputed with zeros or negative ones, as appropriate. (h) Integration with the KEGG database refines the data to 2144 common genes. (i) Clinical dataset preprocessing involves dropping or imputing missing values, leading to *M* common samples for multi-omic analysis.

Furthermore, the reference datasets (clinical features, Ensembl dataset, gene regulatory network) were collected and integrated with processed multi-omics dataset for providing labels, gene annotation and knowledge graph construction). The common samples/patients across multi-omics datasets and concatenated clinical dataset were intersected, leaving *M* common samples, where *M* = 138 if all of the multi-omics datasets were utilized (see **Figure 8A** and **Table 5**).

**Figure 8.**
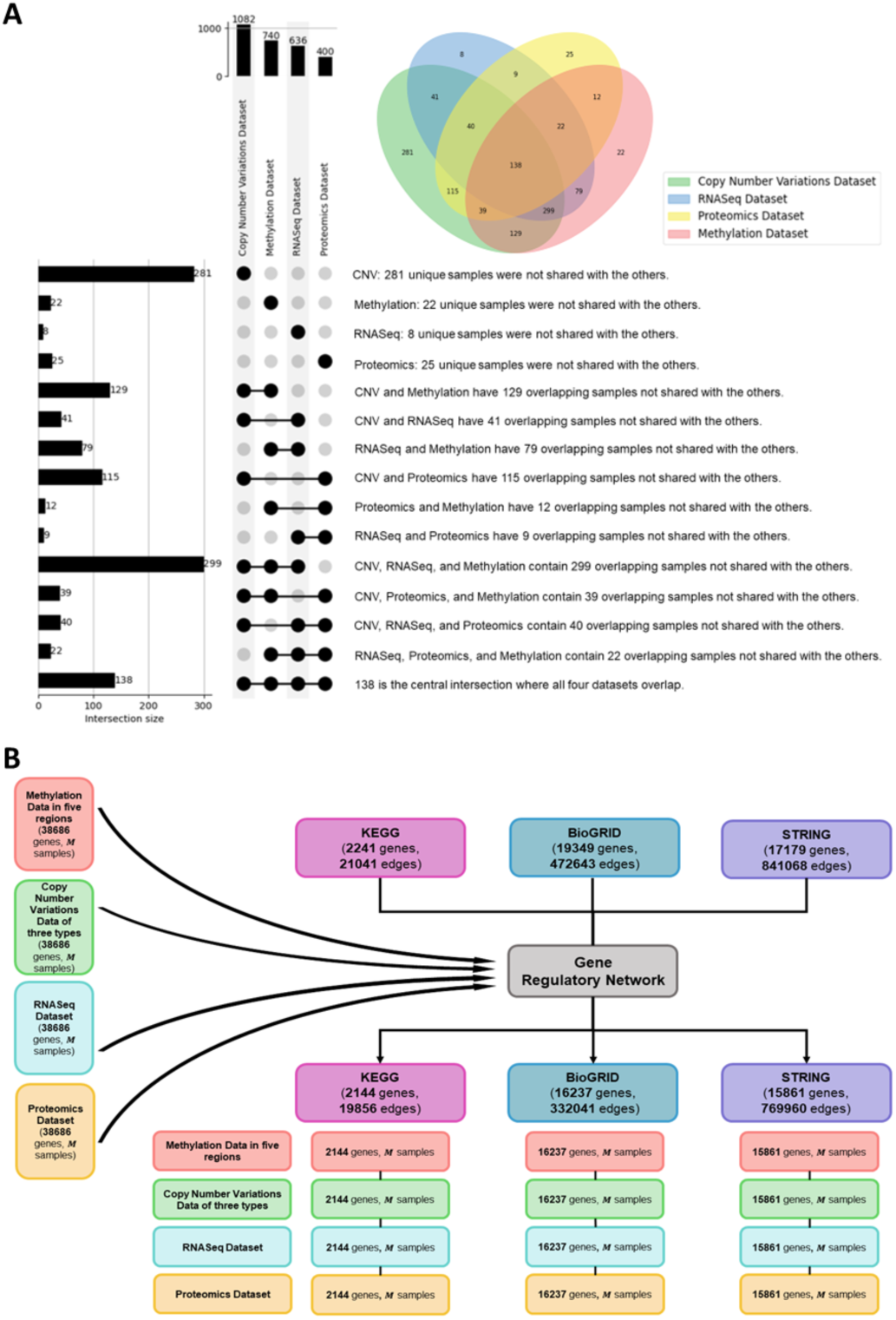
Details for intersection for samples gene regulatory network. (a)The intersection of omics data with clinical samples, finally *M* samples across all omics data. (b) The intersection of the multi-omic data with different options of gene regulatory network

**Table 5.**
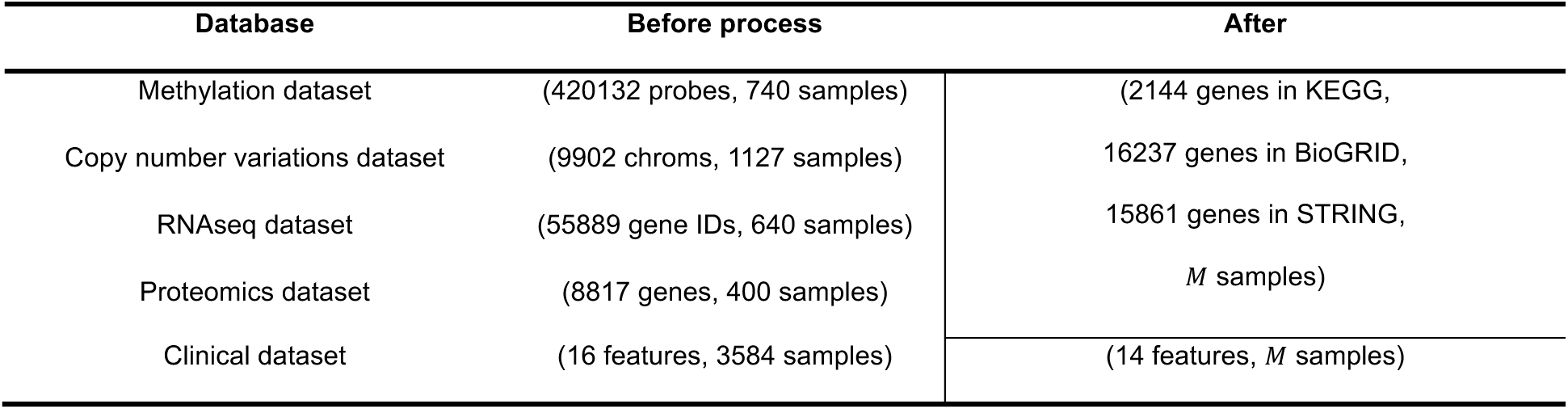
Dimensions of the ROSMAP multi-omics data before and after the data processing.

Additionally, the Ensembl dataset was utilized to annotate genes in multi-omics data, which led to the following dimensions: methylation dataset (17606 genes, *M* samples), Copy Number Variations dataset (6419 genes, *M* samples), RNASeq dataset (37898 genes, *M* samples), and proteomics dataset (7817 genes, *M* samples). The union of genes from the multi-omics datasets was achieved by aggregating all unique genes across the datasets, yielding a total of 38,686 genes. Subsequently, genes for constructing the knowledge graph were selected through the intersection of multi-omics datasets with gene regulatory networks—specifically, (KEGG^24,40^ (2241 genes, 21041 edges), BioGRID^23,41^ (19349 genes, 472643 edges) and STRING^42^ (17179 genes, 841068 edges)), resulting the number of gene entities as 2144, 16237 and 15861 (see **Figure 8B**).

## Discussion

Our study introduced mosGraphGen, a novel computational tool that enables the generation of multi-omics signaling graphs for each sample by mapping the multi-omics data onto a biologically relevant multi-level signaling network. With mosGraphGen, AI model developers can seamlessly apply and assess their models using these multi-omics signaling graphs. We conducted evaluations of mosGraphGen using multi-omics datasets from both cancer and Alzheimer’s disease (AD) samples. Furthermore, we have made the code, examples, and tutorials for mosGraphGen open-source and publicly accessible for the research community via GitHub: https://github.com/FuhaiLiAiLab/mosGraphGen. The following sections provide the details of the methodology and results. Users can build the mosGraphs using the two widely used multi-omics datasets following the instructions for their own down-stream analysis.

Given the exploratory nature of this study, it is important to acknowledge the inherent limitations that present opportunities for in-depth exploration in subsequent research endeavors. For instance, there is potential for enhancing the biological significance of the background signaling network to improve the biological relevance of multi-omics data integration. It is also important to biologically or experimentally validate these mosGraphs in specific studies, which can further indicate the reliability and utility, and further improve the model. Furthermore, we aspire to develop an interactive tool that allows users to generate mosGraphs without the need for coding, particularly tailored for well-known and widely used multi-omics datasets and data platforms. There are a few potential future improvements and extensions of mosGraphGen model. For example, the nucleic acid sequence can be included to indicate the specific and fine-level genetic mutations and epigenetic changes. It will increase the computational cost dramatically. Also, the metabolomic data is critical to understand disease pathogenesis. Thus, it is also important to integrate metabolomic data using the known-protein metabolites interactions or their co-expressions to identify the synergistic effects of the multi-omic and metabolomic features.

Moreover, the single cell omic datasets have been generated to understand disease molecular mechanisms at single cell level and potential cell-cell interactions. Thus, it is interesting and important to expand the mosGraphs for single-cell omics data. However, it’s important to note that dealing with a substantially larger number of mosGraphs for single-cell multi-omics data, given the presence of tens of thousands of single cells in individual studies, poses a challenge in terms of data storage and analysis. In addition to the numeric multi-omic data, the known knowledge or descriptive/text information of individual proteins, like the annotation and description of the functions of individual proteins, can be added. The large language models (LLMs) can be used to convert the description/text information into numeric feature to be combined with the numeric omic features for the target and signaling pathway inference/mining from the mosGraphs. Therefore, novel graph AI models that can analyze the large-scale mosGraphs and identify the essential disease associated targets and signaling pathways are needed.

## Acknowledgement

This study was partially supported by NIA R56AG065352 (to Li), 1R21AG078799-01A1 (to Li/Province), 1RM1NS132962-01 (to Dickson/Marco/Cooper/Li), 1R01LM013902-01A1 (to Li).

## References

1. Grossman RL, Heath AP, Ferretti V, et al. Toward a Shared Vision for Cancer Genomic Data. New England Journal of Medicine. 2016;375(12):1109–1112. doi:10.1056/nejmp1607591

2. Sanchez-Vega F, Mina M, Armenia J, et al. Oncogenic Signaling Pathways in The Cancer Genome Atlas. Cell. 2018;173(2):321–337.e10. doi:10.1016/j.cell.2018.03.035

3. Goldman MJ, Craft B, Hastie M, et al. Visualizing and interpreting cancer genomics data via the Xena platform. Nat Biotechnol. 2020;38(6):675–678. doi:10.1038/s41587-020-0546-8

4. Barretina J, Caponigro G, Stransky N, et al. The Cancer Cell Line Encyclopedia enables predictive modelling of anticancer drug sensitivity. Nature. 2012;483(7391):603–607. doi:10.1038/nature11003

5. De Jager PL, Ma Y, McCabe C, et al. A multi-omic atlas of the human frontal cortex for aging and Alzheimer’s disease research. Sci Data. 2018;5(1):1–13.

6. Bennett DA, Buchman AS, Boyle PA, Barnes LL, Wilson RS, Schneider JA. Religious orders study and rush memory and aging project. Journal of Alzheimer’s disease. 2018;64(s1):S161–S189.

7. Allen M, Carrasquillo MM, Funk C, et al. Human whole genome genotype and transcriptome data for Alzheimer’s and other neurodegenerative diseases. Sci Data. 2016;3. doi:10.1038/sdata.2016.89

8. Neff RA, Wang M, Vatansever S, et al. Molecular Subtyping of Alzheimer’s Disease Using RNA Sequencing Data Reveals Novel Mechanisms and Targets. Vol 7.; 2021. https://www.science.org

9. Li F, Eteleeb A, Buchser W, et al. Weakly activated core inflammation pathways were identified as a central signaling mechanism contributing to the chronic neurodegeneration in Alzheimer’s disease. bioRxiv. Published online 2021.

10. Greenwood AK, Montgomery KS, Kauer N, et al. The AD Knowledge Portal: A Repository for Multi-Omic Data on Alzheimer’s Disease and Aging. Curr Protoc Hum Genet. 2020;108(1). doi:10.1002/cphg.105

11. Raghavachari N, Wilmot B, Dutta C. Optimizing Translational Research for Exceptional Health and Life Span: A Systematic Narrative of Studies to Identify Translatable Therapeutic Target(s) for Exceptional Health Span in Humans. Journals of Gerontology - Series A Biological Sciences and Medical Sciences. 2022;77(11):2272–2280. doi:10.1093/gerona/glac065

12. Hao N, Li Y, Jiang Y, et al. Evolutionary-Conserved, Molecular Mechanisms Can Simul-Taneously Influence All Causes of Death. WADDINGTON’S LANDSCAPE OF CELL AGING DISRUPTION OF CPG ISLAND-MEDIATED CHROMATIN ARCHITECTURE AND TRANSCRIPTIONAL HOMEOSTASIS DURING AGING SESSION 1120 (SYMPOSIUM) THE LONGEVITY CONSORTIUM: MULTI-OMICS INTEGRATIVE APPROACH TO DISCOVERING HEALTHY AGING AND LONGEVITY DETERMINANTS Chair.

13. Deelen J, Evans DS, Arking DE, et al. A meta-analysis of genome-wide association studies identifies multiple longevity genes. Nat Commun. 2019;10(1). doi:10.1038/s41467-019-11558-2

14. Lee J, Hyeon DY, Hwang D. Single-cell multiomics: technologies and data analysis methods. Exp Mol Med. 2020;52(9):1428–1442.

15. Baysoy A, Bai Z, Satija R, Fan R. The technological landscape and applications of single-cell multi-omics. Nat Rev Mol Cell Biol. 2023;24(10):695–713. doi:10.1038/s41580-023-00615-w

16. Subramanian I, Verma S, Kumar S, Jere A, Anamika K. Multi-omics data integration, interpretation, and its application. Bioinform Biol Insights. 2020;14:1177932219899051.

17. Sedgewick AJ, Benz SC, Rabizadeh S, Soon-Shiong P, Vaske CJ. Learning subgroup-specific regulatory interactions and regulator independence with PARADIGM. Bioinformatics. 2013;29(13):i62–i70.

18. Yu MK, Ma J, Fisher J, Kreisberg JF, Raphael BJ, Ideker T. Visible Machine Learning for Biomedicine. Cell. 2018;173(7):1562–1565. doi:10.1016/j.cell.2018.05.056

19. Ma J, Yu MK, Fong S, et al. Using deep learning to model the hierarchical structure and function of a cell. Nat Methods. 2018;15(4):290–298. doi:10.1038/nmeth.4627

20. Kuenzi BM, Park J, Fong SH, et al. Predicting Drug Response and Synergy Using a Deep Learning Model of Human Cancer Cells. Cancer Cell. 2020;38(5):672–684.e6. doi:10.1016/j.ccell.2020.09.014

21. van den Berg R, Kipf TN, Welling M. Graph convolutional matrix completion. ArXiv. Published online 2017.

22. Szklarczyk D, Franceschini A, Wyder S, et al. STRING v10: protein–protein interaction networks, integrated over the tree of life. Nucleic Acids Res. 2015;43(D1):D447–D452.

23. Oughtred R, Stark C, Breitkreutz BJ, et al. The BioGRID interaction database: 2019 update. Nucleic Acids Res. 2019;47(D1):D529–D541.

24. Kanehisa MGS, Goto S. KEGG: kyoto Encyclopedia of Genes and Genomes. Nucleic Acids Res. 2000;28:27–30. doi:10.1093/nar/28.1.27

25. Slenter DN, Kutmon M, Hanspers K, et al. WikiPathways: a multifaceted pathway database bridging metabolomics to other omics research. Nucleic Acids Res. 2018;46(D1):D661–D667.

26. Zhang H, Chen Y, Payne P, Li F. Mining signaling flow to interpret mechanisms of synergy of drug combinations using deep graph neural networks. doi:10.1101/2021.03.25.437003

27. Dong Z, Zhang H, Chen Y, Payne PRO, Li F. Interpreting the Mechanism of Synergism for Drug Combinations Using Attention-Based Hierarchical Graph Pooling. Cancers (Basel). 2023;15(17). doi:10.3390/cancers15174210

28. Zhang H, Goedegebuure SP, Ding L, et al. M3NetFlow: a novel multi-scale multi-hop multi-omics graph AI model for omics data integration and interpretation. doi:10.1101/2023.06.15.545130

29. Feng J, Province M, Li G, Payne PRO, Chen Y, Li F. PathFinder: a novel graph transformer model to infer multi-cell intra- and inter-cellular signaling pathways and communications. doi:10.1101/2024.01.13.575534

30. Li F, Dong Z, Payne P, et al. Highly accurate disease diagnosis and highly reproducible biomarker identiication with PathFormer. Published online 2023. doi:10.21203/rs.3.rs-3576068/v1

31. Wang T, Shao W, Huang Z, et al. MOGONET integrates multi-omics data using graph convolutional networks allowing patient classification and biomarker identification. Nat Commun. 2021;12(1):3445.

32. Li X, Ma J, Leng L, et al. MoGCN: a multi-omics integration method based on graph convolutional network for cancer subtype analysis. Front Genet. 2022;13:806842.

33. Gao H, Zhang B, Liu L, Li S, Gao X, Yu B. A universal framework for single-cell multi-omics data integration with graph convolutional networks. Brief Bioinform. 2023;24(3):bbad081.

34. Rajadhyaksha N, Chitkara A. Graph Contrastive Learning for Multi-omics Data. arXiv preprint arXiv:230102242. Published online 2023.

35. Butler JEF, Kadonaga JT. The RNA polymerase II core promoter: A key component in the regulation of gene expression. Genes Dev. 2002;16(20):2583–2592. doi:10.1101/gad.1026202

36. Saxonov S, Berg P, Brutlag DL. A Genome-Wide Analysis of CpG Dinucleotides in the Human Genome Distinguishes Two Distinct Classes of Promoters.; 2005. www.pnas.orgcgidoi10.1073pnas.0510310103

37. Saintenac C, Jiang D, Akhunov ED. Targeted analysis of nucleotide and copy number variation by exon capture in allotetraploid wheat genome. Genome Biol. 2011;12(9). doi:10.1186/gb-2011-12-9-r88

38. Duttke SHC, Lacadie SA, Ibrahim MM, et al. Human promoters are intrinsically directional. Mol Cell. 2015;57(4):674–684. doi:10.1016/j.molcel.2014.12.029

39. Vanaja A, Yella VR. Delineation of the DNA Structural Features of Eukaryotic Core Promoter Classes. ACS Omega. 2022;7(7):5657–5669. doi:10.1021/acsomega.1c04603

40. Ogata H, Goto S, Sato K, Fujibuchi W, Bono H, Kanehisa M. KEGG: Kyoto encyclopedia of genes and genomes. Nucleic Acids Res. Published online 1999:28. doi:10.1093/nar/27.1.29

41. Stark C, Breitkreutz BJ, Reguly T, Boucher L, Breitkreutz A, Tyers M. BioGRID: a general repository for interaction datasets. Nucleic Acids Res. 2006;34(suppl_1):D535–D539.

42. Szklarczyk D, Gable AL, Lyon D, et al. STRING v11: protein–protein association networks with increased coverage, supporting functional discovery in genome-wide experimental datasets. Nucleic Acids Res. 2019;47(D1):D607–D613.

